# Time elapsed between choices in a probabilistic task correlates with repeating the same decision

**DOI:** 10.1101/643965

**Authors:** Judyta Jabłońska, Łukasz Szumiec, Piotr Zieliński, Jan Rodriguez Parkitna

## Abstract

Reinforcement learning causes an action that yields a positive outcome more likely to be taken in the future. Here, we investigate how the time elapsed from an action affects subsequent decisions. Groups of C57BL6/J mice were housed in IntelliCages with access to water and chow *ad libitum*; they also had access to bottles with a reward: saccharin solution, alcohol or a mixture of the two. The probability of receiving a reward in two of the cage corners changed between 0.9 and 0.3 every 48 h over a period of ~33 days. As expected, in most animals, the odds of repeating a corner choice were increased if that choice was previously rewarded. Interestingly, the time elapsed from the previous choice also influenced the probability of repeating the choice, and this effect was independent of previous outcome. Behavioral data were fitted to a series of reinforcement learning models. Best fits were achieved when the reward prediction update was coupled with separate learning rates from positive and negative outcomes and additionally a “fictitious” update of the expected value of the nonselected choice. Additional inclusion of a time-dependent decay of the expected values improved the fit marginally in some cases.

## Introduction

Positive reinforcement increases the probability of repeating actions that were previously rewarded. When a choice between two alternatives is offered, the one with greater expected value is more likely to be taken. If the potential outcomes change due to shifts in the environment, sampling the available choices and balancing the exploitation and exploration of these choices become necessary. These behaviors are controlled by the brain’s reward system, which signals prediction error and performs updates of expectation from any action or cue contingencies (Schultz, 2015). The strategy employed by humans, e.g., (Holroyd & Coles, 2002; O’Doherty *et al*., 2003), and animals, e.g., (Fiorillo *et al*., 2003; Bayer & Glimcher, 2005), when faced with a choice between probabilistic rewards is consistent with reinforcement models that rely on temporal difference learning and may be predicted with algorithms developed in the machine learning field (Sutton *et al*., 2018). Reinforcement learning plays an essential role in adaptive behavior, and impaired decision making has been a major focus in research on the etiology of neuropsychiatric disorders (Maia & Frank, 2011).

Thus far, experimental models used in studies of reinforcement learning have been based on choices made in environments with minimized distractions, short time scales and large numbers of choices performed in quick succession (e.g., (Clark *et al*., 2004; Izquierdo *et al*., 2017)). The mechanisms controlling intervals between responses have received considerable attention, with a focus on the ability to select optimal times for maximizing rewards under paradigms where a specific delay in response after a cue was required for optimum result or with different cues signaling various lengths of delay to reward (Gibbon, 1977; Killeen & Fetterman, 1988; Fiorillo *et al*., 2008; Gershman *et al*., 2014; ligaya *et al*., 2018). Dopamine signaling in the striatum was attributed to an essential role in the control of the response delay, integrating reinforcement learning and timing of responses (Daw *et al*., 2006; Ludvig *et al*., 2008). It should be noted, however, that response timing is likely influenced by multiple pathways associated with the reward system, and recent reports show that serotonergic neurons may play a particularly significant role (Hinton & Meck, 1997; Matias *et al*., 2017; Iigaya *et al*., 2018). While the mechanisms controlling interval timing affect behaviors on the scale from seconds to hours, due to methodological limitations, behavioral models are mostly focused on intervals shorter than a minute. As a consequence, memory decay or other processes occurring on longer scales are rarely considered (e.g., (Greggers & Menzel, 1993; Collins & Frank, 2012; Collins *et al*., 2017)). The focus on short intervals and exclusion of any confounding influences limits variability, conforms with some of the underlying assumptions in reinforcement learning models (e.g., resembles a Markov process), and has the major advantage of allowing for correlation of behavior with neuronal spiking activity. However, whether uninterrupted sequences of quick decisions are an adequate approximation of reinforcement learning under normal environmental conditions is arguable.

Here, we assess reinforcement in a probabilistic choice reversal learning paradigm in which mice were not compelled to perform the task in any way, and choices were performed freely over a period of weeks. We tested two different types of primary rewards: saccharin and alcohol solutions, which differ mechanistically in the way they affect the reward system. We found that in a large number of cases, choices were significantly influenced by previous outcomes; however, the interval between choices played a comparable, if not greater, role.

## Methods

### Animals

Experiments were performed on female C57BL/6J mice bred at the Maj Institute of Pharmacology of the Polish Academy of Sciences in Krakow. Mice were housed in a conventional facility in Plexiglas cages (Type II L, 2–5 animals per cage) with aspen laboratory bedding (MIDI LTE E-002, Abedd) and nest building material. Breeding rooms had a 12 h light/dark cycle, with an ambient temperature of 22 ± 2°C and humidity of 40-60%. Animals were provided with a piece of aspen wood for chewing after weaning. Mice had *ad libitum* access to water and chow (RM1 A (P), Special Diets Services). All experiments were conducted in accordance with the European Union guidelines for the care and use of laboratory animals (2010/63/EU). Experimental protocols were reviewed and approved by the II Local Bioethics Committee in Krakow (permits 1000/2012 and 1159/2015). Behavior was tested on female mice to reduce the risk of aggressive behaviors. The experimental groups are summarized in Table 1.

**Table 1.** Experimental groups.

| Cohort | n | Reward | Starting age | Mean initial weight ± SEM [g] |
| --- | --- | --- | --- | --- |
| I | 14 | Alcohol 4% (w/v) | 8 weeks | 18.79 ± 1.64 |
| I | 14 | Saccharin 0.1% (w/v) | 8 weeks | 18.62 ± 1.79 |
| II | 14 | Alcohol 4% (w/v) | 8 weeks | 19.04 ± 2.36 |
| II | 14* | Alcohol+Saccharin | 8 weeks | 19.59 ± 1.95 |
| III | 14 | Saccharin 0.1% (w/v) | 8 weeks | 19.49 ± 2.25 |
| IV | 14 | Water (control) | 8 weeks | 17.96 ± 0.19 |

\*One animal lost its transponder and was excluded.

### Probabilistic choice task

The IntelliCage apparatus (New Behavior, Switzerland) has a base made of transparent plastic (55 x 37.5 x 20.5 cm) with a metal cover and custom corner compartments. Each of the cage corners is a small chamber that houses two 250 ml bottles, with nozzles accessible through guillotine doors (Figure 1A). The size of the corner allows only one animal to enter the corner and access the bottles. Before being introduced to the IntelliCage, mice are implanted with radio frequency identification chips (RFID chips, UNO PICO ID, AnimaLab, Poland). An antenna inside the corner detects the chip and reports the animal number to the controlling software, which triggers preprogrammed events. The cage recorded the following parameters: temperature and luminosity in 1 minute intervals, presence of an animal in a corner (combined reading from a thermal sensor and an RFID antenna), crossing of photocell beams placed in the doors leading to the bottles, and lickometer contacts (the animal closing a circuit between the floor grating and the metal dipper of the bottle). Experiments were performed on groups of 14 female mice per cage. The number of animals was based on our previous experiments (Smutek *et al*., 2014; Ruud *et al*., 2019). At the start of the test, mice were introduced to the IntelliCage, which had standard bedding and contained 4 plastic “houses” that the animals used as nests to sleep during the day. The environmental conditions during the experiment were the same as those in the breeding rooms, with food and water available *ad libitum*. Exact schedules for each experiment are shown in Figure 1B. At the start of the experiments, all corners had bottles filled with water. When a mouse entered a corner and the RFID chip was detected, both guillotine doors blocking access to the bottles would open with a 0.5-s delay. The doors were closed when the mouse left the corner or 10 s after a lick of a bottle was detected. The initial period lasted between 4 and 7 days, and the mice were monitored daily to check whether all of them had learned to drink from the bottles. Then, the adaptation stage started, and a reward (saccharin 0.1% (w/v), alcohol 4% (w/v), a mixture of the two or plain water) became available in two of the corners. We chose a low concentration of alcohol to ensure high preference and to limit the effects of inebriation on learning. Saccharin was selected over sacharose to exclude nongustatory effects and to avoid clogging of the dippers. The adaptation stage lasted ~3 weeks, and bottle positions were changed regularly to reduce the formation of corner preferences (see Figure 1B). Finally, during the main stage of the experiment, the probability of reward access varied between 90% and 30%, with a 2-s delay from an entrance to a corner to the opening of the guillotine doors. Additionally, yellow LED lights in the reward corners were switched on when the animal was detected and switched off if the animal left the corner or after 2 s (irrespective of whether reward access was granted). The LED lights were intended as an additional cue of a choice being in progress to reduce the effect of variability caused by the animal detection mechanism. The positions of the corners with reward bottles were constant, while the probabilities changed between all possible states—90%:30%, 30%:90%, 90%:90% and 30%:30%—as shown in Figure 1.

**Figure 1.**
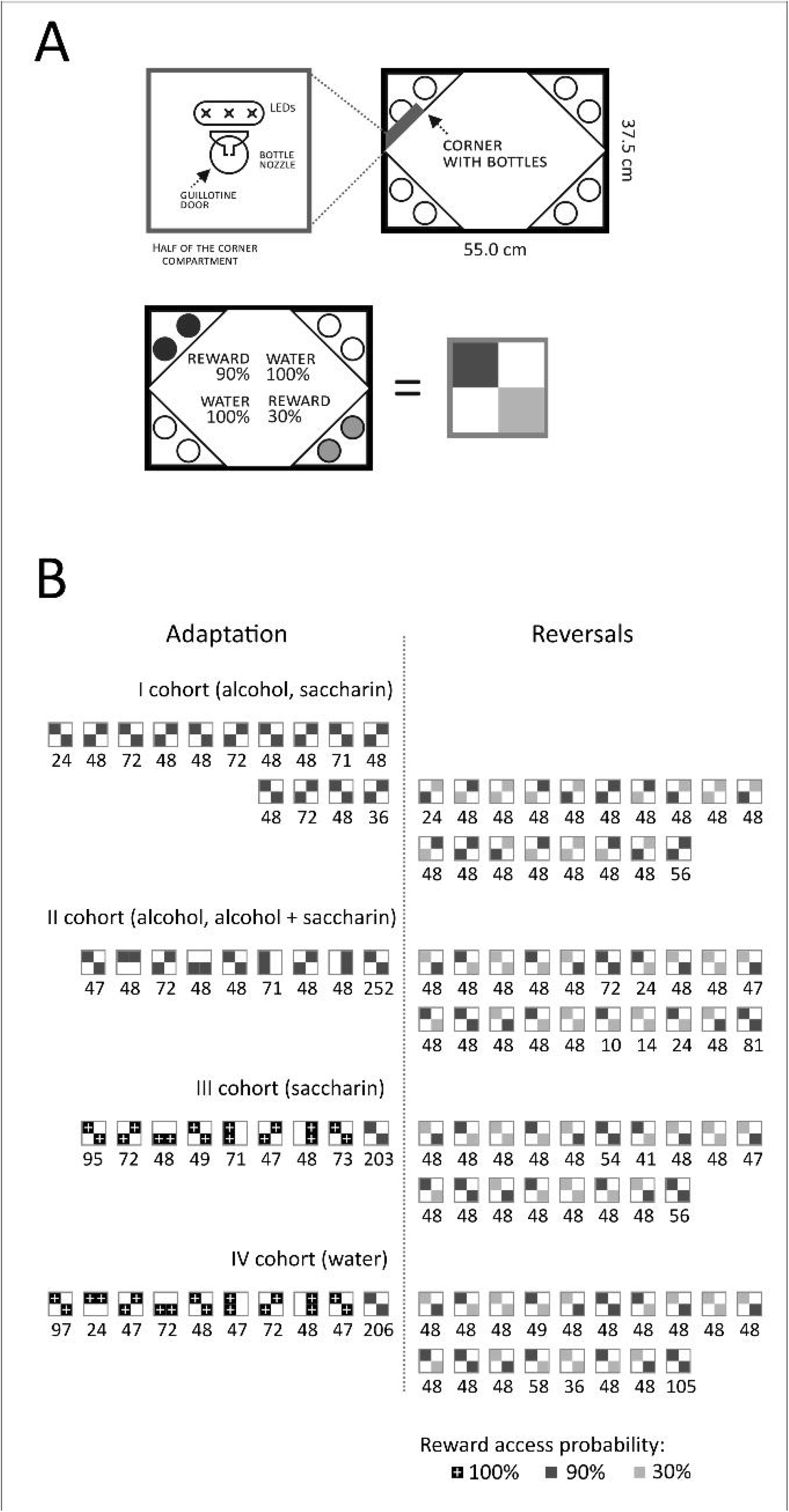
Schematic representation of the IntelliCage and experimental schedules. (A) The diagrams show the basic features of the cage and the guillotine doors through which bottles are accessed. The diagram on the bottom shows an example of the key used to label reward probabilities. (B) For each of the groups indicated on the left, the phases are represented with white, grey or black boxes. Black boxes with white lines indicate corners with full reward access (100%), dark grey boxes indicate the high probability of reward access (90%), light grey boxes indicate the low probability (30%) and white indicates access to water. The duration of each phase in hours is presented below the corresponding boxes.

### Models

Logistic regression was used to assess the effects of the outcome and time elapsed from the previous choice on the odds of behavior in terms of repeating (“stay”) the previous decision. The following formula was used:

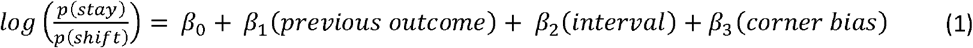

where *β*_0_ is the intercept (preference independent of predictors), *β*_1_ represents the outcome of the previous choice (binary, equals 1 after win), *β*_2_ is the effect of the time interval elapsed from the previous choice (per minute of interval), and *β*_3_ is the corner (binary). The Wald test was used to assess the significance of the predictors. Data corresponding to two animals were excluded due to extreme corner preference, with only 2/135 and 1/650 alternative choices, respectively.

The following reinforcement-learning models were tested. First was the random choice model, which assumed that the chance of either choice was equal at each step. The second model was the “noisy win-stay-lose-shift” model described by Wilson and Collins (Wilson & Collins, 2019). Briefly, the model predicts “stay” after a rewarded attempt and “shift” when no reward is received, adding a probability of random choice defined by the parameter *ϵ*,

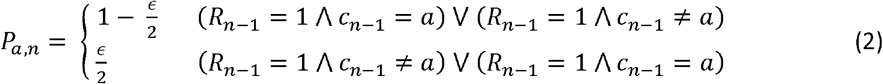

where *P_a,n_* is the probability of selecting corner *a* at choice *n, c*_*n*−1_ is the previous choice, and *R*_*n*−1_ is the value of the reward after previous choice (1 or 0). Further models fitted were based on Q-learning (Watkins & Dayan, 1992; Sutton *et al*., 2018), starting with the most “basic”:

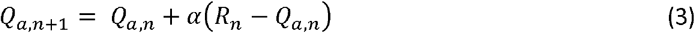

where *Q_a,n_* represents the expected value of selecting corner *a* at step *n, R_n_* is the reward received at step *n* (1 or 0), and *α* is the learning rate. The model was coupled with a softmax policy,

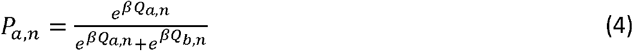

where *P*_*a,n*+1_ is the probability of choosing corner *a* at step *n*, and *β* is an “inverse temperature” parameter that determines the extent to which the difference between expected rewards affects choice. Probabilities were limited to a minimum of 0.001 and maximum of 0.999 to limit the effect of extreme values on the log likelihood sum. The first modification to the model was “dual” learning rates, which depended on the reward value:

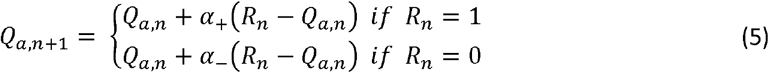

where *α*_+_ and *α*_−_ are separate learning rates for rewarded and nonrewarded choices. The next modification included an update for expected values for both the choice taken (*Q_a,n_*) and the nonselected alternative (*Q_b,n_*), partly based on the “fictitious” update described previously (Hampton *et al*., 2007):

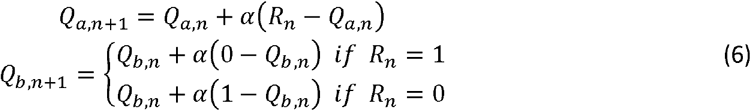

An extension of this approach is the “hybrid” model, which combines (5) and (6) (Cieślak *et al*., 2018):

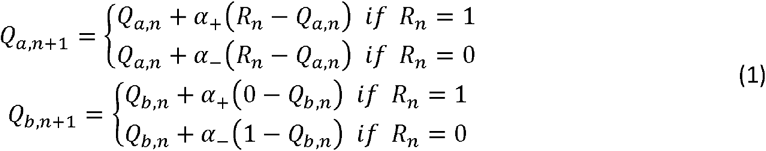

Nest we tested a group of models that include a parameter representing the effects of memory performance, starting with the “forgetful” model (Collins & Frank, 2012), where equation (3) was followed by:

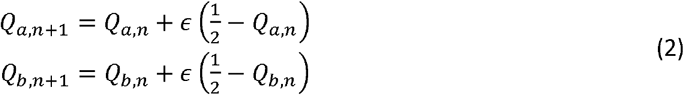

where *ϵ* (in the range from 0 to 2) represents the limited ability to recall the expected values. To introduce the effect of the interval between choices, we tested models incorporating time-dependent exponential decay of the expected value (“Qd”) or the inverse temperature parameter in the policy (“*β*d”). The exponential decay component was based on observations on memory decay in humans (Murre & Dros, 2015) and previous theoretical considerations (Woźniak *et al*., 1995). The interval was defined as the time elapsed since the last choice (end of the corner visit to start of the current visit). The starting model introduced equal decay of expected values for both choices:

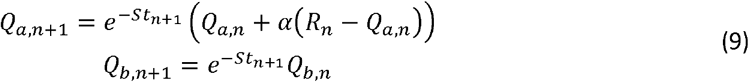

where *t*_*n*+1_ is the time interval between choices *n* and *n* + 1, and *S* is a parameter representing memory performance (“storage”). An *S* value of 0 would indicate no memory decay. Due to computational limits, the maximum interval length was set to 660 minutes. Two extensions of the model were considered. The first is a fictitious update (“Qd+fictitious”):

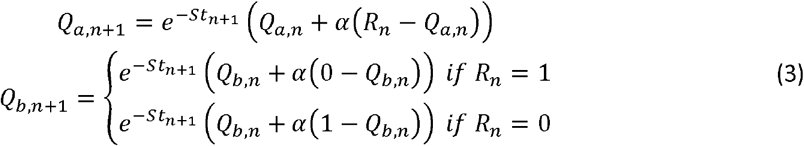

Second, separate learning rates depending on the previous outcome (“Qd+hybrid”):

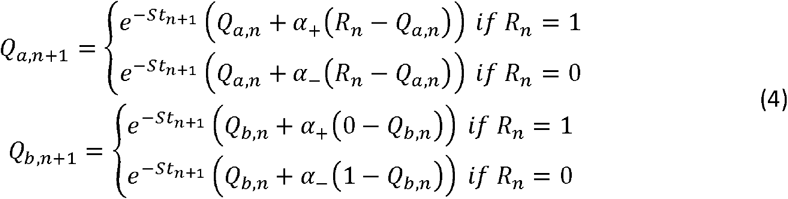

As an alternative way to account for the effects of the time intervals, we considered a decay of the inverse temperature parameter in the policy (“*β*d”):

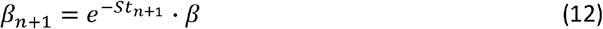

where *t*_*n*+1_ is the interval from the previous choice and *β*_*n*+1_ replaces *β* in equation (4). The final model (“*β*d-ficitious”) combined equations (6) and (12). The optimal parameters for each model were selected based on the lowest sum of the negative logarithms of likelihood:

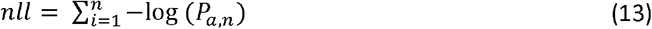

where *P_a,n_* is the probability of the actual choice predicted by the model. Models with more than one parameter (all except noisy win-stay-lose-shift) were fitted using the Nelder and Mead simplex method (Nelder & Mead, 1965) implemented in the R *optim* function using all possible combinations of the following starting points: {0.05, 0.15, 0.35, 0.55, 0.75, 0.95} in the cases of *α* and *S*, {0.05, 0.35, 0.75, 1, 1.25, 1.35, 2} in the case of *ϵ*, and {0.25, 1.25, 2.25, 3.25, 4.25} in the case of *β*. Parameter values were limited to the ranges of (0, 1) for *α* and *S*, (0, 2) for *ϵ* and (0, 50) in the case of *β*. In the noisy win-stay-lose-shift model, the optimal parameter value was assessed by calculating nil for *ϵ* in the range of 0.001 to 1.999 in 0.001 steps. Model fits were compared using Akaike’s information criterion (Akaike, 1974):

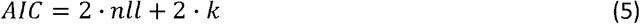

where *k* is the number of free parameters in the model. Δ*AIC* values were calculated by subtracting the AIC value corresponding to the “basic” model from the results obtained for each of the other models.

### Data analysis and statistics

All analyses were performed using R (R Core Team, 2017), and the scripts used to analyze the data are available at https://github.com/jmjablons/model-inteli-research2019. Statistical significance was assessed using the Kruskal-Wallis test followed *a posteriori* by the Dunn test adjusted for multiple comparisons using the Benjamini-Hochberg correction. One-sample and paired-sample comparisons were performed with the Wilcoxon test. The significance (α) level was set at 5%. The complete set of behavioral data is available at https://figshare.com/s/4d39377b1ce5cc3c24c2.

## Results

### An unconstrained probabilistic choice task

We tested the behavior of group-housed female mice that could freely select between two probabilistic reward alternatives over an extended period of time (Figure 1). Three types of rewards were offered in separate experiments: a saccharin solution (0.1% w/v), alcohol (4% w/v), and a mixture of the two (alcohol+saccharin). As a control, in a separate experiment, the reward was replaced with plain tap water. All experiments were split into two stages. The first was adaptation, during which the positions of the rewards were switched to reduce potential biases towards cage corners and to allow for the development of an alcohol preference. During the second, main stage, the positions of the reward bottles were fixed, but the probabilities of opening access to the bottles changed. Additionally, irrespective of stage, the animals also had access to water bottles in the two remaining corners of the cage (empty squares in Figure 1B). As shown in Figure 2, the activity of animals, i.e., the total number of visits to the corners during each 48-h period, was similar for all types of rewards and remained stable throughout the main stage of testing. The total time the two reward corners were occupied (i.e., the cage detected the presence of a mouse) was generally shorter than 4 h per 48-h period (Supplement Figure 1). The only exception was in one of the “alcohol” cohorts, where mice were detected for extended periods of time in the reward corners over three 48-h periods. Based on these results, we assume that competition for corner access had no appreciable effect on choices, which is consistent with the results we have reported previously (Smutek *et al*., 2014).

**Figure 2.**
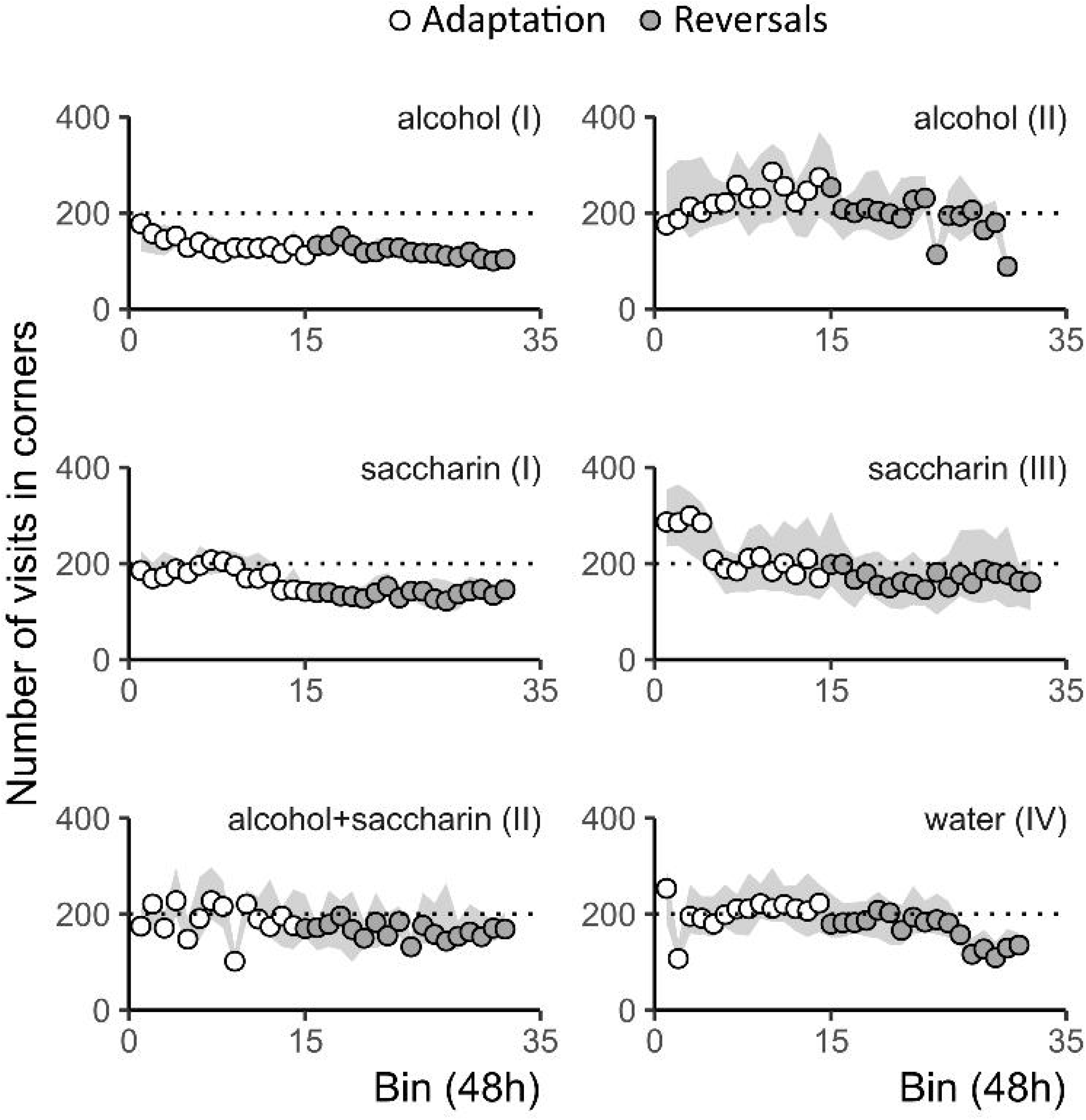
Animal activity in the IntelliCages. The graphs show the median total number of visits in all corners of the cage per 48 h. Circles correspond to activity during the adaptation, and black points correspond to probability reversals. The ribbon shows 1st and 3rd quartiles. Each graph summarizes the results from one group of animals (Table 1). The type of reward used in the experiment is indicated above.

The majority of animals showed preference for the reward, calculated as the number of licks on the dippers of the bottles containing rewards, divided by the total number of licks on all bottles during the final 96 h of the adaptation stage (Figure 3A). The median preferences were 79.4% for saccharin, 80.3% for the saccharin-alcohol mixture, and 65.0% for alcohol (all three significantly higher than 50%, Figure 3A). Not all the animals showed preference; there were 3 and 7 mice in the saccharin and alcohol groups, respectively, that were recorded to have less than 50% of all licks on reward bottles. No significant preference was observed in the water control group, where the median was 42.2%. Saccharin-containing solutions were significantly more preferred than both water or alcohol, and alcohol was significantly more preferred than water (Figure 3A; Kruskal-Wallis test H = 31.94 p = 5.39 × 10^-7^). Despite the preferences of rewards over water, the median fractions of all visits in the reward corners during the stage with probability reversals remained approximately 0.5 (Supplement Figure 2). These data show that animals explored and sampled bottles in all corners over the course of the entire experiment.

**Figure 3.**
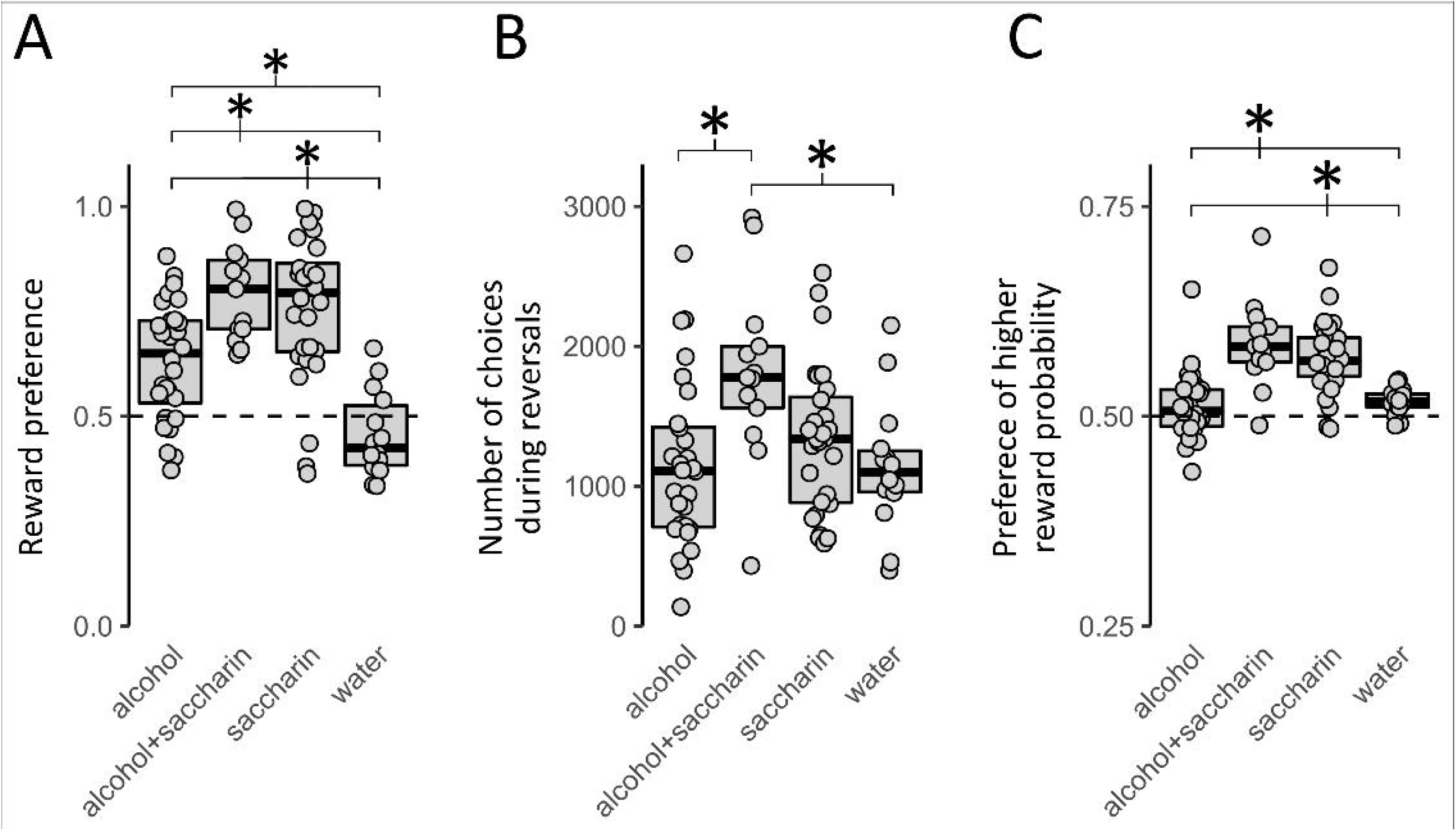
Basic summary of choices. (A) Reward preference during the last 96 h before the start of the reversals stage. (B) Number of attempts (visits longer than 2 s in corners with the reward) at the reversals stage. (C) Fraction of attempts corresponding to higher reward probability. The type of reward is indicated below the graphs. Each dot represents a single animal. Boxplots show medians, 1st and 3rd quartiles. Significant differences between the medians are indicated with stars (* indicates p < 0.05, Dunn’s post hoc test with Benjamini-Hochberg correction).

Further analyses were performed only on visits in the reward corners that lasted sufficiently long for the outcome to occur (>2 s) - designated “choices”. There was considerable individual variation in the numbers of choices performed; the minimum was 137 (an animal in the alcohol group), and the maximum was 2932 (a mouse in the alcohol+saccharin group). The median number of choices was significantly higher in the alcohol+saccharin group than in the alcohol group (Figure 3B, Kruskal-Wallis test, H = 11.413, p = 9.692 × 10^-3^). During the probability reversal stage of the experiment, the median preference for the alternative with a higher reward probability was 56.6% for saccharin and 58.3% for the alcohol+saccharin mixture, compared to 50.5% and 51.7% for alcohol and water, respectively (Figure 3C). In all these cases, except the alcohol-treated group, the median preference was significantly higher than random. Moreover, the preference of larger reward probability was higher in the case of saccharin-containing solutions compared to both alcohol or water (Kruskal-Wallis, H = 35.121, p = 1.149 × 10^-7^). The small but significantly greater than random preference for water may suggest that the opening of the guillotine doors became a conditioned reinforcer.

### Factors affecting choice

First, we assessed the frequency with which the animals repeated a previously rewarded choice (“win-stay”) or shifted to the alternative when no reward was obtained (“lose-shift”). As shown in Figure 4, the frequency of “win-stay” depended on the interval between attempts and was significantly larger at intervals between choices longer than 10 minutes compared to shorter than 2 minutes. This was the case for alcohol (Figure 4A), alcohol+saccharin (Figure 4B) and saccharin (Figure 4C) but not when water was offered instead (Figure 4D). The same was observed in the cases of choices after a “lose”, at longer intervals, the probability of “shift” was decreased in the cases of all rewards (Figure 4A-C), but not in the water control, where in fact an opposite effect was present (higher probability of “shift” at longer intervals, Figure 4D).

**Figure 4.**
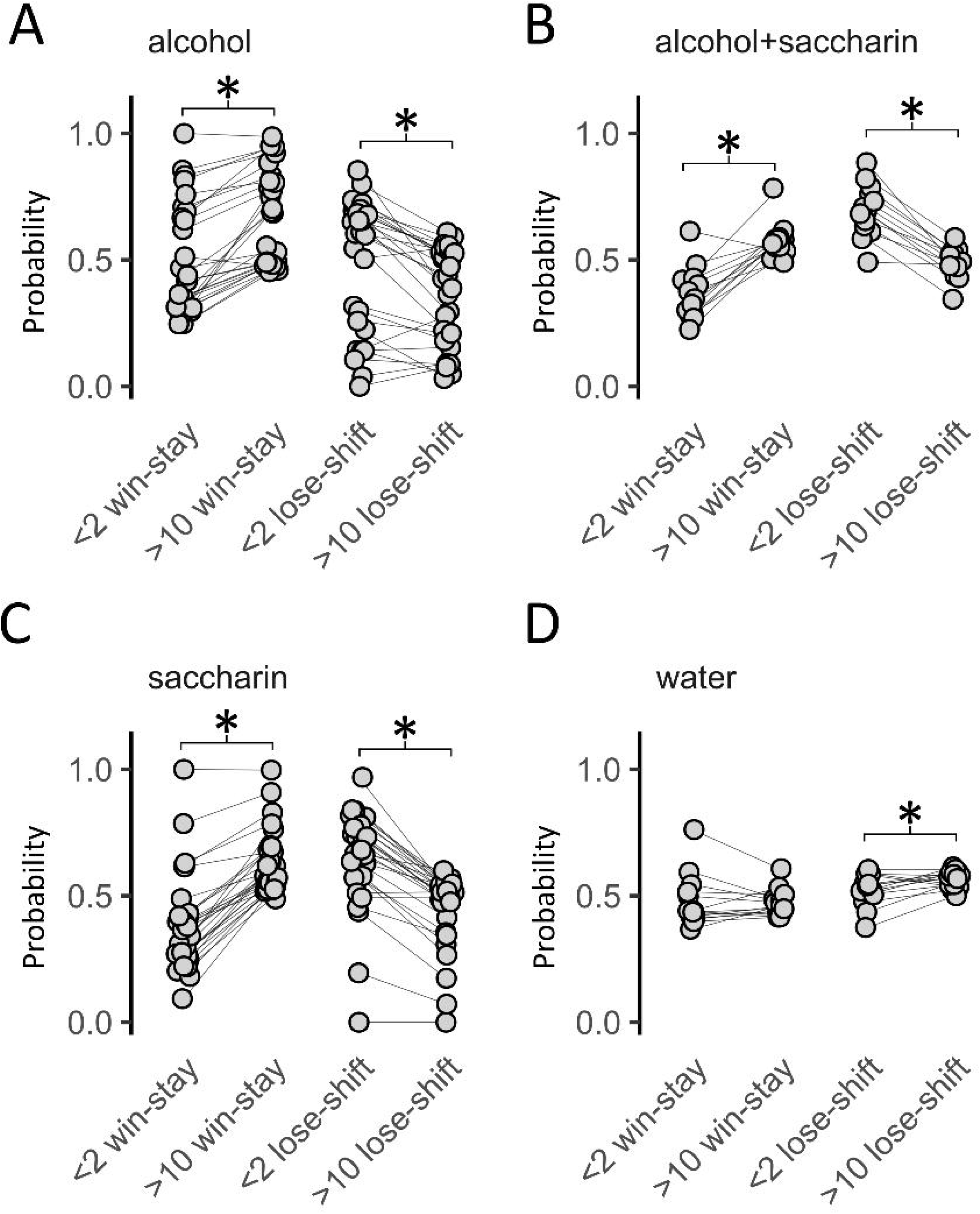
Fractions of “win-stay” and “lose-shift” reactions divided by length of interval between attempts (shorter than 2 minutes and longer than 10 minutes). Each dot represents a single animal. Significant differences between the medians are indicated with stars (* indicates p< 0.05, Wilcoxon test for paired samples).

To further assess the effect of interval and previous reward on choice, we used logistic regression with three predictors: corner bias, outcome of the previous attempt (“win” or “lose”), and time interval from the previous attempt. Examples of the regression curves are presented in Figure 5. The first example (Figure 5A) corresponds to a mouse from the saccharin group. The “stay” behavior was more frequent at longer intervals, whereas “shifts” were more frequent for shorter intervals. This effect is apparent in the distribution of the time intervals. Conversely, in the example drawn from the water control group, no effects of interval or previous outcome are apparent (Figure 5B). Accordingly, the logistic regression shows that the probability of a “stay” decision increased with the length of the interval, and the effect of previous outcome is noticeable as a shift of the curve (“win” vs. “lose). A complete summary of logistic regression analyses for all mice is shown in Figure 6A-D. In the majority of cases (63/81), the models indicate a significant inherent propensity towards “stay” or “shift” responses independent of the predictors considered (the regression intercept, Figure 6A). Animals often had significant corner bias (68/81), with varied individual corner preferences (Figure 6D). As anticipated, in most cases, the model also indicated a significant effect of the previous outcome: 45/81, or excluding the water control group and the interval length: 45/81 and 36/67, respectively (Figure 6B, C). When only significant predictors were considered, the median values of the odds ratios were similar, and no significant effects of reward type were observed. In summary, more than half of the mice that were offered alcohol or saccharin solutions were significantly more likely to “stay” after a “win” and were also more likely to choose “stay” when more time had elapsed from the previous choice. It should be noted, however, that the significant effect of the intercept may indicate that the model has limited accuracy.

**Figure 5.**
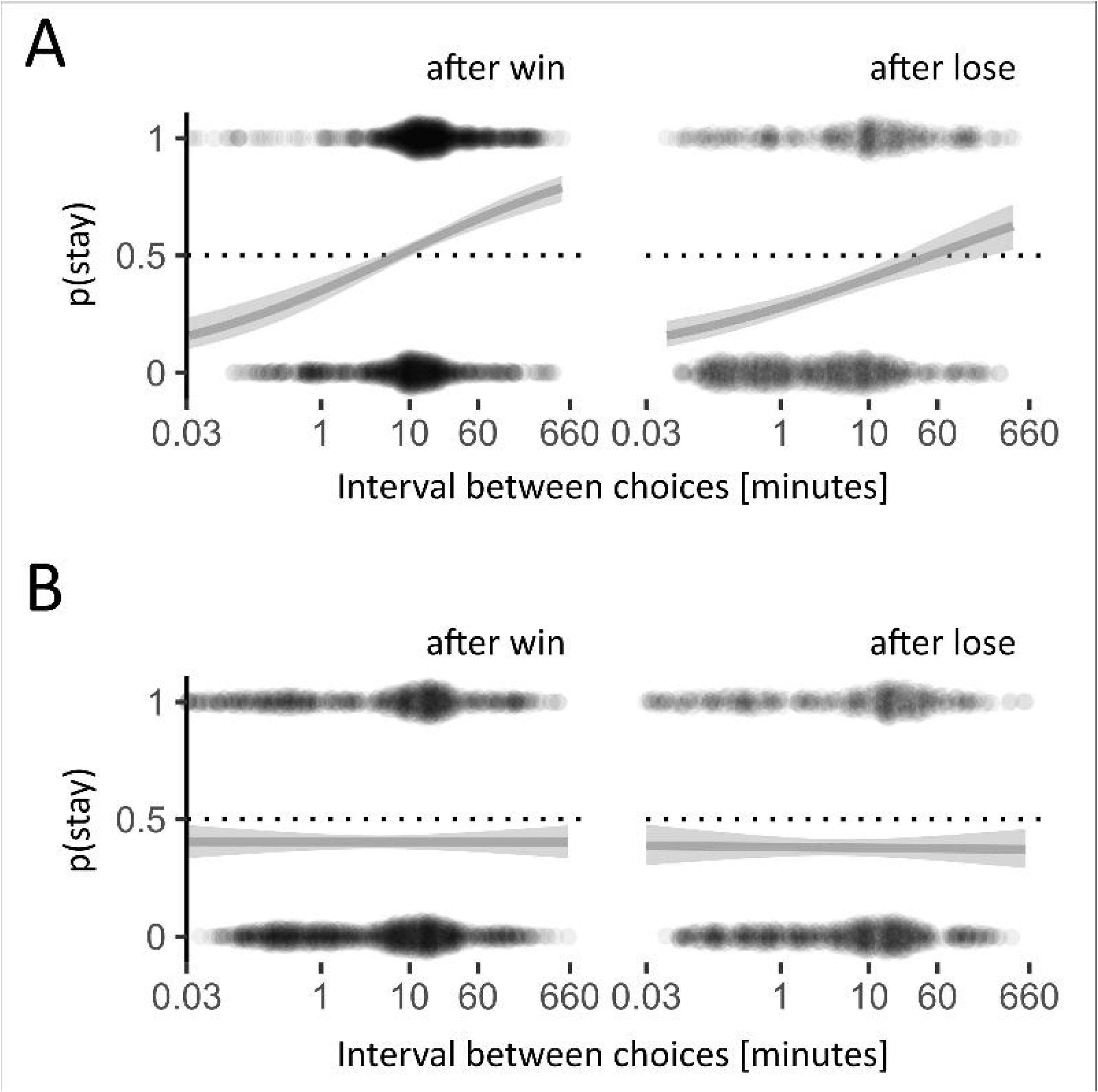
Examples of behavior of a single animal from the saccharin (A) and water (B) groups, respectively. The line is the fitted logistic regression of the probability of “stay” depending on the previous outcome and interval. The dots above and below the curve show raw results used for the regression (probability equal to 1 denotes “stay”, 0 is “shift”).

**Figure 6.**
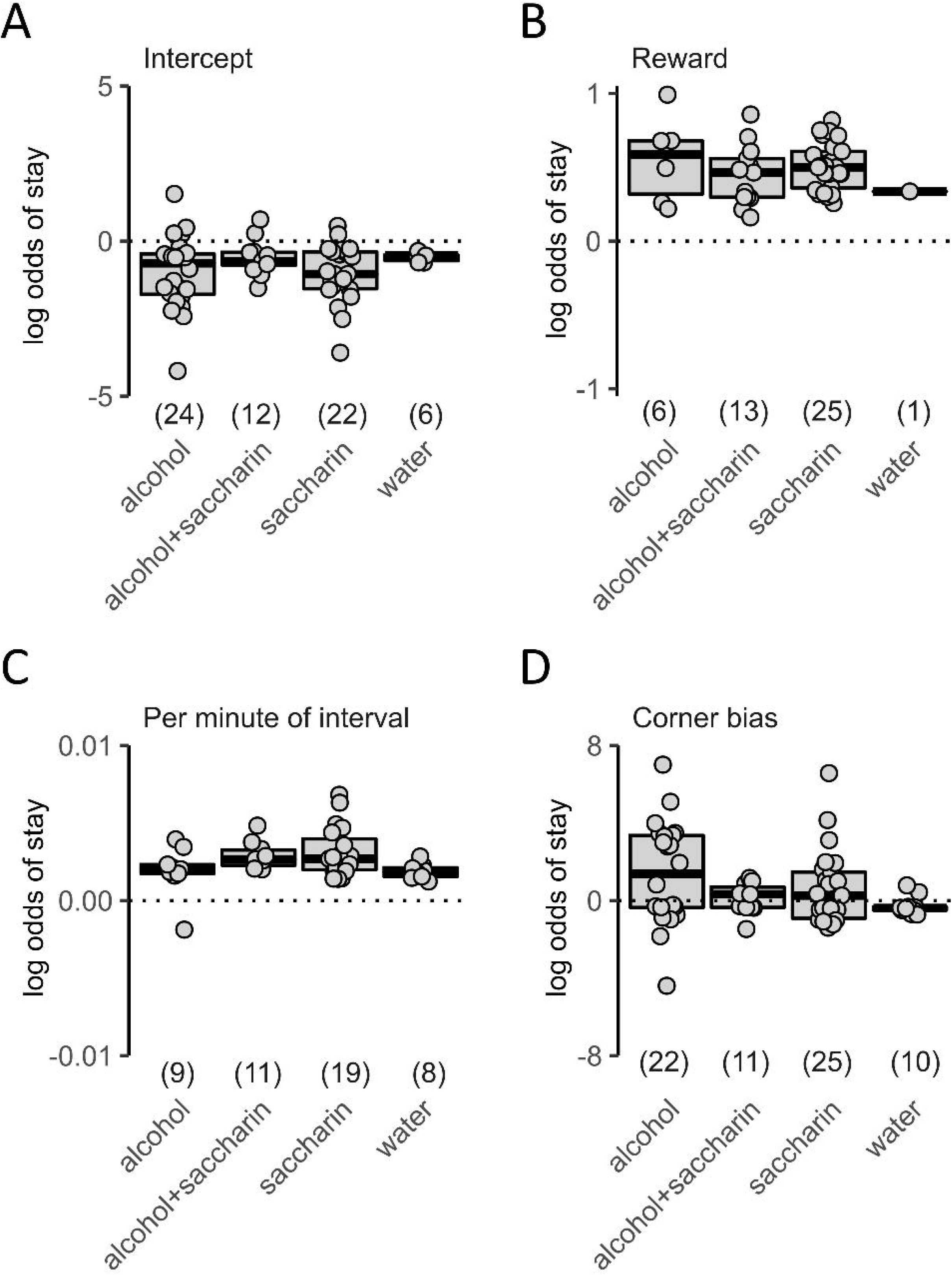
Effects of previous outcome and time interval on choice. Graphs show the change in the log odds of “stay” for each of the predictors: (A) the intercept independent of predictors, (B) after a rewarded choice (“win”), (C) as a function of the interval, and (D) due to inherent corner bias, respectively. Each dot represents a single animal, and only significant results are shown (Wald test P < 0.05). Boxplots show medians, 1st and 3rd quartiles. The type of reward and the number of cases where the predictor was significant are indicated below the graphs.

### Learning models

The preference for the choice associated with a higher reward probability and significant effect of previous outcome on choice imply reinforcement learning. Thus, we fitted various learning models to determine which model assumptions yielded the closest match to the observed behavior. First, we considered a model that selects “stay” after “win” and “shift” after “lose”, with an additional chance that the choice is random instead (“noisy win-stay”, (Wilson & Collins, 2019)). Second, we tested a group of models based on the assumption that the expected value of a choice is updated based on the observed prediction error (Rescorla & Wagner, 1972; Watkins & Dayan, 1992; Sutton *et al*., 2018). The simplest, “basic”, model assumes a single learning rate and updates the expected value. The “dual” model had separate learning rates for negative and positive values of the prediction error. The “fictitious” model updates the value of both options simultaneously (Hampton *et al*., 2007), and “hybrid” added two learning rates to the fictive update (Cieślak *et al*., 2018). Then, we considered models introducing the effects of memory performance and the length of the interval between choices. The “forgetful” model introduces a component related to reverting to a base state of expected values (controlled by the *ϵ* parameter). The decay models adjust the expected reward (“Qd”) or the inverse temperature parameter (“*β*d”) depending on the length of the interval. In both cases, we have also considered the effect of a fictitious update (“+fictitious” or “+hybrid”). In addition to the types of models listed, a “random” choice rule with every choice probability equal to 0.5 was added as a negative control.

Models were fitted by finding the lowest sum of the negative log likelihoods (nll). We used Akaike’s information criterion (AIC) to assess the goodness of fit. A summary of AIC score differences (Δ*AIC*) is shown in Figure 7, the results for best models are summarized in Table 2, and all individual values are provided in Supplementary Table 1. Overall, the models approximated the observed behavior better than a purely random choice approach for all rewards except the water control. Moreover, there were larger differences in the goodness of fit for rewards that produced stronger preference (i.e., saccharin and alcohol+saccharin). The best scoring models, “hybrid” and “Qd+hybrid”, shared two features: separate learning rates for positive and negative outcomes and a fictitious update of the expected value of the nonselected alternative. Modeled positive and negative learning rates differed (for all rewards except water), with median *α*_−_ values close to 0, while median *α*_+_ values were in the range of 0.017 to 0.05 (Table 2). Optimal values of *β* in the hybrid models ranged from 0.81 to 2.73. Median AIC differences between the two top scoring models were approximately 2 or lower, which suggests similar goodness of fit. This is not unexpected since the “Qd+hybrid” model is equivalent to “hybrid” when *S* = 0. Accordingly, the median optimal values of *S* were very low and corresponded to an ~1% loss of the expected value per hour in the alcohol or saccharin groups (Table 2). We note that in the cases where the “Qd+hybrid” hybrid model had the best goodness of fit, the values tended to be above the median.

**Figure 7.**
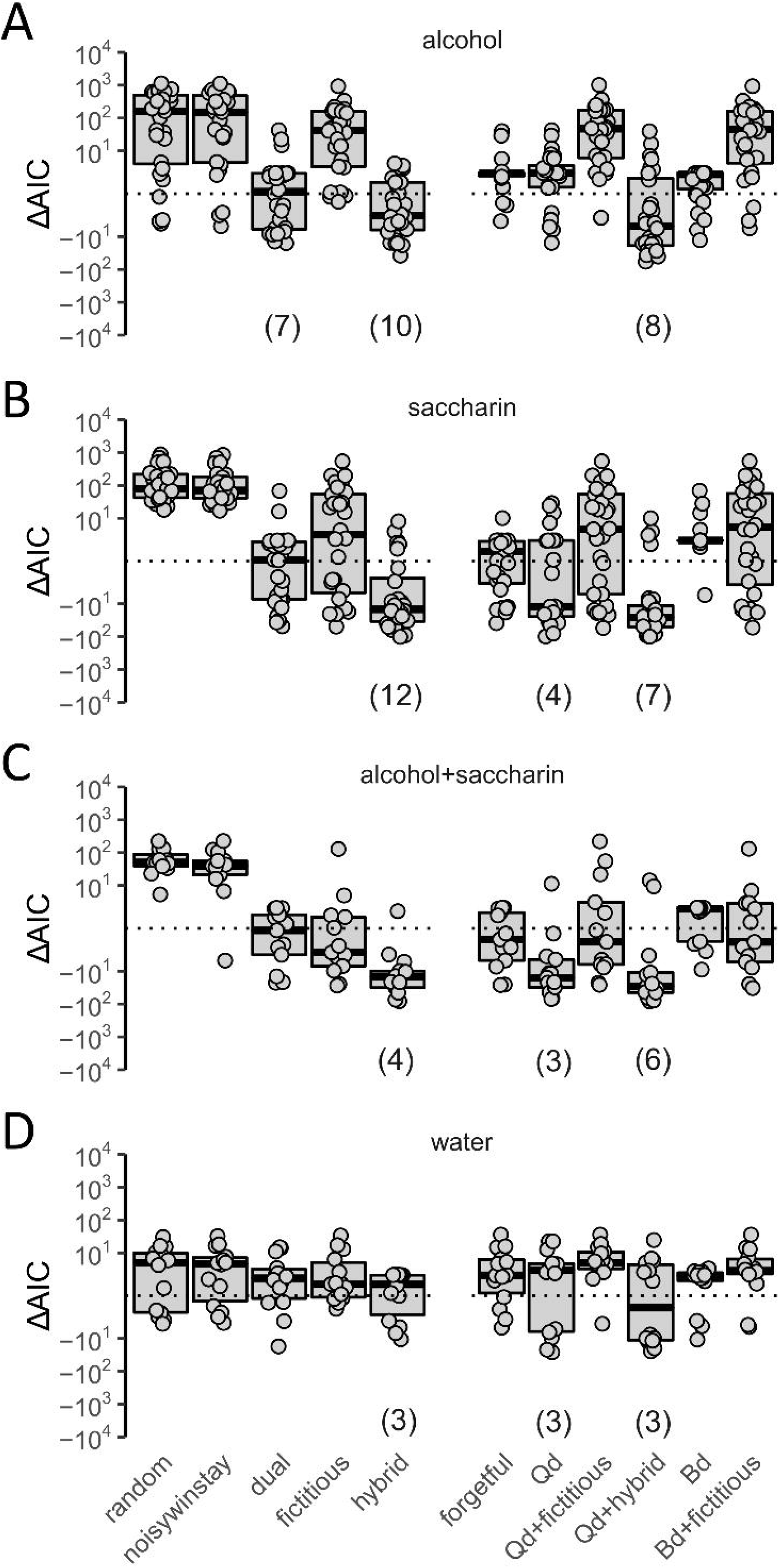
Comparison of reinforcement learning model fits based on Δ*AIC* values. Panels correspond to (A) alcohol, (B) saccharin, (C) alcohol+saccharin and (D) water. Each point represents an individual mouse, the boxplots show medians, 1st and 3rd quartiles. The numbers in parentheses shown below plots indicate how many times the lowest AIC score was achieved for that model, for the best three models.

**Table 2.** Summary of optimized models’ parameters.

|  | alcohol | saccharin | alcohol+saccharin | water |
| --- | --- | --- | --- | --- |
| **basic** |  |  |  |  |
| $\alpha$ | 0.00601*<br>(0.00426-0.0112) | 0.00912<br>(0.00634-0.0139) | 0.11<br>(0.0331-0.173) | 0.00179<br>(0.00016-0.0731) |
| $\beta$ | 6.2<br>(3.32-7.52) | 3.85<br>(3.22-4.97) | 0.956<br>(0.682-3.33) | 5.05<br>(0.312-50) |
| **hybrid** |  |  |  |  |
| $\alpha_+$ | 0.024<br>(0.0113-0.0676) | 0.035<br>(0.024-0.0513) | 0.0877<br>(0.0298-0.104) | 0.00193<br>(0.0001-0.0396) |
| $\alpha_-$ | $4.14 \times 10^{-9}$<br>($4.1 \times 10^{-10}$-$3.4 \times 10^{-4}$) | $4.71 \times 10^{-9}$<br>($1.46 \times 10^{-9}$-$1.21 \times 10^{-8}$) | $2.65 \times 10^{-8}$<br>($3.27 \times 10^{-10}$-0.0026) | $4.61 \times 10^{-5}$<br>($1.14 \times 10^{-8}$-0.623) |
| $\beta$ | 2.54<br>(1.62-3.1) | 1.64<br>(1.31-1.8) | 0.899<br>(0.795-1.25) | 1.22<br>(0.215-36.6) |
| **Qd hybrid** |  |  |  |  |
| $\alpha_+$ | 0.017<br>($1.03 \times 10^{-6}$-0.031) | 0.0303<br>(0.0179-0.0427) | 0.0499<br>(0.0156-0.0971) | 0.00392<br>($1.10 \times 10^{-7}$-0.017) |
| $\alpha_-$ | $3.61 \times 10^{-5}$<br>($2.43 \times 10^{-8}$-0.94) | $3.02 \times 10^{-8}$<br>($9.07 \times 10^{-9}$-0.00348) | $3.39 \times 10^{-8}$<br>($1.14 \times 10^{-8}$-$8.86 \times 10^{-6}$) | $8.40 \times 10^{-5}$<br>($1.69 \times 10^{-8}$-1) |
| $\beta$ | 2.73<br>(1.07-3.52) | 1.89<br>(1.54-2.35) | 1.62<br>(1.01-1.92) | 0.813<br>(0.447-13.9) |
| $S$ | $9.16 \times 10^{-5}$<br>($1.97 \times 10^{-5}$-0.116) | $7.82 \times 10^{-5}$<br>($1.56 \times 10^{-7}$-0.000512) | 0.000783<br>(0.00052-0.0511) | 0.0332<br>(0.00715-0.534) |

\*Median value; numbers in parentheses below the median represent 5%-95% confidence intervals calculated using bootstrapping.

## Discussion

We show reinforcement learning in mice under conditions where there is no enforced schedule and the actions being reinforced are not being compelled by food or drink deprivation. The animals remained in a group and could behave freely in the cage environment throughout the entire procedure. The only cost of a choice was a short wait period (2 s) before the outcome was presented, and the costs of missed opportunities were theoretically negligible. There was no trial and session structure and no maximum number of potential rewards. In approximately half of the animals tested, when a choice was rewarded, the probability of repeating it was significantly increased as assessed using a logistic regression model. The observed effect of previous outcomes is weaker than that reported in studies employing classical models (e.g., (Frank *et al*., 2004; Kwak *et al*., 2014; Cieślak *et al*., 2018)). No mouse exceeded 75% in their preference for the corner with higher reward probability, and median preference values were lower than 60%. Intuitively, the simplest explanation for the low preference for the higher reward probability would be lack of missed opportunity cost and hence limited advantage of performing optimal choices. The correlation between the number of choices and the preference for the higher probability of reward was −0.3, - 0.72, 0.045 and −0.35 for alcohol, alcohol+saccharin, saccharin and water, respectively (Pearson’s r). While the value in the case of alcohol+saccharin is significant, this appears to be an exception, and we do not think that this gives evidence that animals compensated for a smaller fraction of rewarded choices by increasing the number of attempts. Furthermore, while both median preferences of the higher reward probability and the analysis of individual behavior confirm reinforcement learning in a large fraction of the mice tested, nevertheless, it should be noted that animals did not strictly try to maximize the number of rewards obtained. Mice generally made fewer than 100 choices per 48 h, compared to 120 or more trials completed within 30 minutes in some Skinner box experiments (e.g., (Cieślak *et al*., 2018)). The limited number of choices is not unexpected considering that in this case, consuming rewards does not satisfy essential needs; however, it poses a problem with regard to defining what constitutes an optimal behavioral strategy in the test.

A major novel observation is the relation between the type of choice made correlated with the interval between decisions. As could be intuitively expected, when a choice was not rewarded, the delay to try again was usually shorter. At the same time, the length of the interval correlated with an increased probability of repeating the same choice, irrespective of the previous outcome. These effects were only observed when a reward was offered, particularly in the case of saccharin-containing solutions and were generally absent in the water control group. It should be stressed that the interval between choices discussed here is not comparable to the interval timing learned when rewards are delivered after a specific delay or different cues predict the length of delay to reward delivery (e.g., (Gershman *et al*., 2014)). Here, the length of the interval had no effect on the size or probability of a reward. Therefore, we assume that the length of the intervals is determined primarily by motivational processes, which would differ for choices performed at short (1-2 minutes) vs. longer (15-20 minutes) intervals. At shorter intervals, the pattern is somewhat similar to spontaneous alternation, which is often observed during exploration of multiarmed mazes (e.g., (Lalonde, 2002)). Conversely, the process that drives choices after longer intervals is intriguing. It does not appear to be optimal, as it promotes choosing “stay” after a nonrewarded choice. Potentially, repeating the choice could serve the purpose of reconsolidating a learned value, though again, the advantage this may offer is unclear. Notably, our finding is limited to female C57BL/6J mice, and whether the effect may be generalized to other species remains unknown. An effect of sex on operant learning in rodents was reported, although it was mostly observed in the context of responses conditioned with aversive stimuli (Dalla & Shors, 2009). In humans, there is ample evidence that women are more risk averse than men (Byrnes *et al*., 1999); however, gender had no significant effects on the win-stay or lose-shift ratios in a probabilistic reversal learning task (den Ouden *et al*., 2013). Assuming that generalization is possible, the findings presented here could, to an extent, explain some of the “irrational” behaviors, i.e., frequent choices of an inferior, lower value alternative.

A preference of higher reward probability implies reinforcement learning; thus, we tested fitting learning models to observed choices. We note, however, that there are important issues to consider with regard to the applicability of reinforcement learning models in this case. First, while the choices in the task have the Markov property, nevertheless, as noted above, the animals did not try to accumulate the maximum number of rewards possible. Moreover, we did not exclude animals from the analysis based on their preference of the higher reward probability or significant effects of previous outcome on choice; thus, in some cases, models were applied to behavior where we have no evidence of reinforcement (e.g., most of the water control group). Despite these limitations, the models approximated behavior better than random choice. The best goodness of fit was achieved when separate learning rates for positive and negative outcomes were applied, and additionally, a fictitious update of the nonselected alternative was also included. The latter is consistent with our previous results in a Skinner box-based model (Cieślak *et al*., 2018); however, we consider it surprising. In the classic report that provided insight into mechanisms responsible for the regret or disappointment over a lost opportunity, information of the outcome for the nonselected choice was provided in a fraction of trials (Coricelli *et al*., 2005). Thus, the subject could at least partly assess the difference in outcome values between the actions. The fictitious model developed by (Hampton *et al*., 2007) assumes that the reward value used applied in the update of the nonselected choice is the opposite of the one obtained, which is similar, but not fully equivalent, to the effect of regret. Here, the cost of lost opportunity is minimal; therefore, an effect of regret does not appear rational. Furthermore, a fictive update implies a more model-based approach to the decision process, which is similar to the conclusion drawn in a study where the optimal strategy involved updating the value of the nonselected option (Huh *et al*., 2009). We also wanted to point out two observations with regard to optimal parameters of the best models. First are the very low learning rates (*α*_+_ and *α*_−_). The median values could be interpreted as no learning from negative outcomes and a very small update after rewarded choices. A larger positive learning rate is consistent with previously reported results (e.g., (Rutledge *et al*., 2009; Cieślak *et al*., 2018)), and the very low values could be speculatively attributed to the negligible cost of choosing the corner with a lower probability of reward. Second, the marginal gain in goodness of models incorporated a time-dependent decay of expected value or the inverse temperature despite the clear correlation between the time interval and the probability of repeating the same choice. Intuitively, this is not surprising, considering that these models fail to predict previous outcome-independent preference for shift choices at short intervals or greater than 0.5 probability of stay choices at very long intervals. The modeled decay constant (*S*) would imply only a minor effect of memory performance, though it should be noted that in the cases where Qd+hybrid had the best goodness, corresponding *S* values were often above median. Nevertheless, based on the results, we would argue that introducing the decay effect does not offer a plausible explanation for the correlation between interval lengths and the probability of “stay” choices.

A second objective of our study was to assess differences in the actions of saccharin and alcohol as reinforcers. The mechanisms by which alcohol and saccharin affect the reward system are inherently different. The effects should be instantaneous and transient in the case of saccharin (signaling from the gustatory system to the midbrain dopamine neurons (Simon *et al*., 2006)) but are delayed by minutes and could be persistent in the case of alcohol (through activation of dopamine release and other mechanisms (Weiss *et al*., 1993; Vengeliene *et al*., 2008)). Nonetheless, drug and natural rewards reportedly produce their long-term effects by acting on the same neuronal circuits (Kelley & Berridge, 2002; Pfarr *et al*., 2018). We should stress that the experiment we report was not intended to model compulsive alcohol drinking, which persists despite increasing cost, reduced value or a risk of negative consequences (Vengeliene *et al*., 2009; Hopf & Lesscher, 2014). Additionally, the concentration of alcohol used was low (4% w/v), and thus blood ethanol levels would be unlikely to match the values achieved in previously described models (Rhodes *et al*., 2005; Rodriguez Parkitna *et al*., 2013). Analysis of the data shows no evidence of impaired learning due to the effects of alcohol on memory performance; the behavior of mice in the saccharin and alcohol+saccharin groups was similar. In both cases, the rewards were highly preferred over water, and there was evidence of reinforcement learning in the majority of animals in those groups. Conversely, while alcohol was preferred over water after the 3-week adaptation stage, only some mice showed significant evidence of reinforcement learning. A possibility that should be considered is that the reward value of the alcohol solution was lower, which could be in line with the reported general preference of sweet taste over drugs in rodents (Ahmed, 2018). A lower reward value compared to saccharin could limit learning based on prediction error, while possibly remaining sufficient to produce a preference over water. Therefore, the only apparent difference in the effects of alcohol and saccharin is that at the concentrations tested, the former was a weaker reinforcer. We cannot exclude the possibility that in a larger tested cohort or when higher alcohol concentrations are offered, a subset of animals would show altered reinforcement learning (as could be hypothesized based on, for instance, (Augier *et al*., 2018)).

In conclusion, the most striking observation emerging from our results is that the effect of time elapsed from an action may affect the probability of repeating it, independent of outcome. Modeling suggests that this effect is not easily explained by memory decay, and it is unclear if it offers an adaptive advantage. We speculate that our observations suggest separate motivation processes driving decisions at short vs. longer intervals; however, further investigation is necessary to verify this hypothesis.

## Acknowledgments

This work was supported by the grant SONATA BIS 2012/07/E/NZ3/01785 from the National Science Centre, Poland, and statutory funds of the Maj Institute of Pharmacology of the Polish Academy of Sciences. JMJ was supported with a PhD stipend from InterDokMed project no. POWR.03.02.00-00-1013/16. We would like to thank Rafał Bogacz for valuable comments and suggestions.

## Author Contributions

JMJ analyzed the data and wrote the paper. ŁS participated in planning and performed the experiments. PZ participated in the data analysis and writing of the paper. JRP planned the experiments, participated in data analysis and wrote the paper.

## Data Accessibility

The complete set of behavioral data collected in this study is available at Figshare https://figshare.com/s/4d39377b1ce5cc3c24c2. The R scripts used for analysis are available at https://github.com/jmjablons/model-inteli-research2019. Should inquiries with regard to the data and analysis tools arise, we will address them to the best of our capacity.

## Abbreviations

AIC: Akaike’s information criterion

---

**Supplementary Figure 1.**
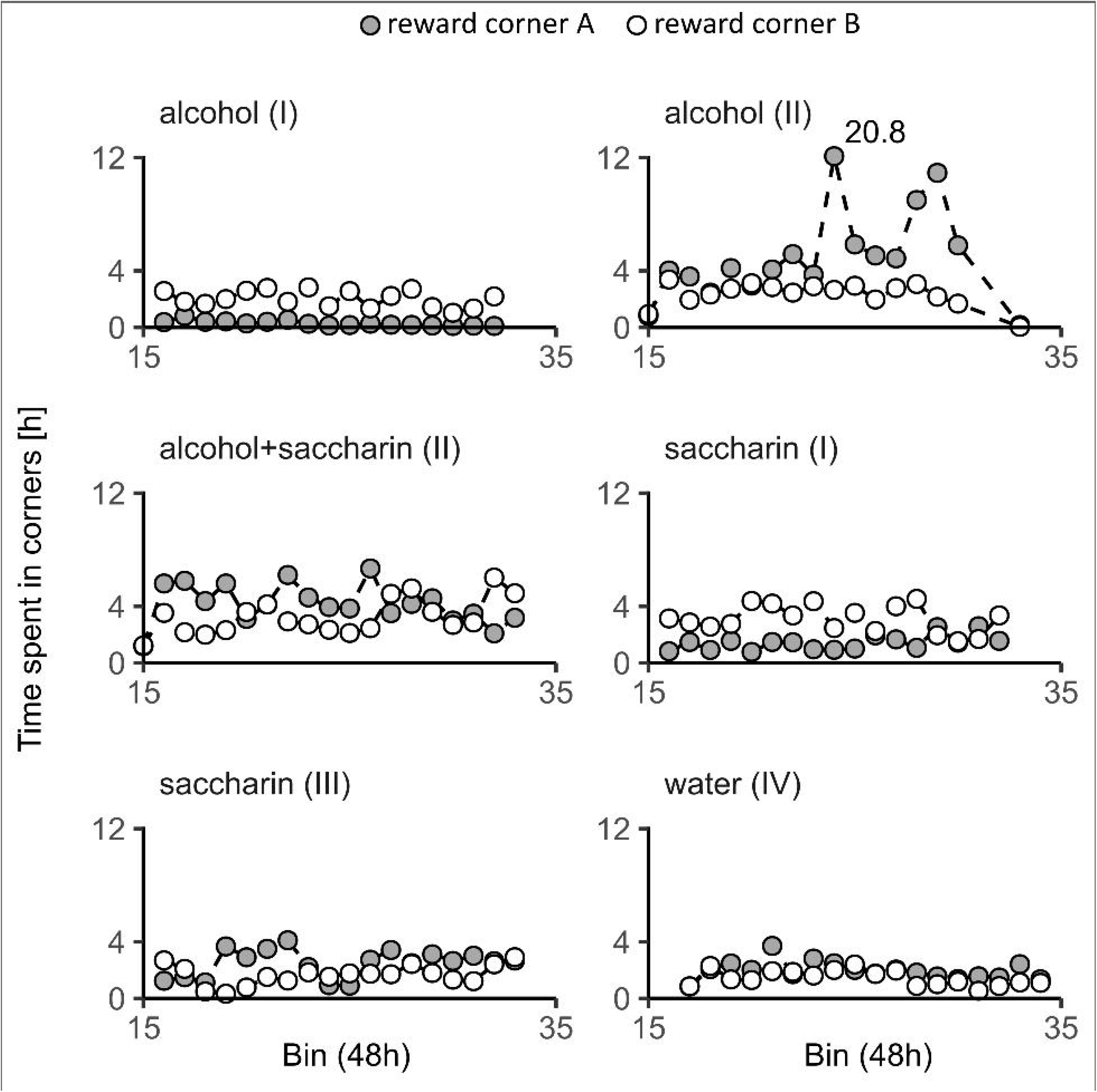
Total number of hours when the cage corners were taken by animals per 48h period. White dots represent the corner with lower internal number (“A”), and grey dots the other (“B”). Each graph summarizes the results from one group of animals tested in one cage. The type of reward offered is indicated above the graphs. Roman numerals in parentheses indicate the animal cohort.

**Supplementary Figure 2.**
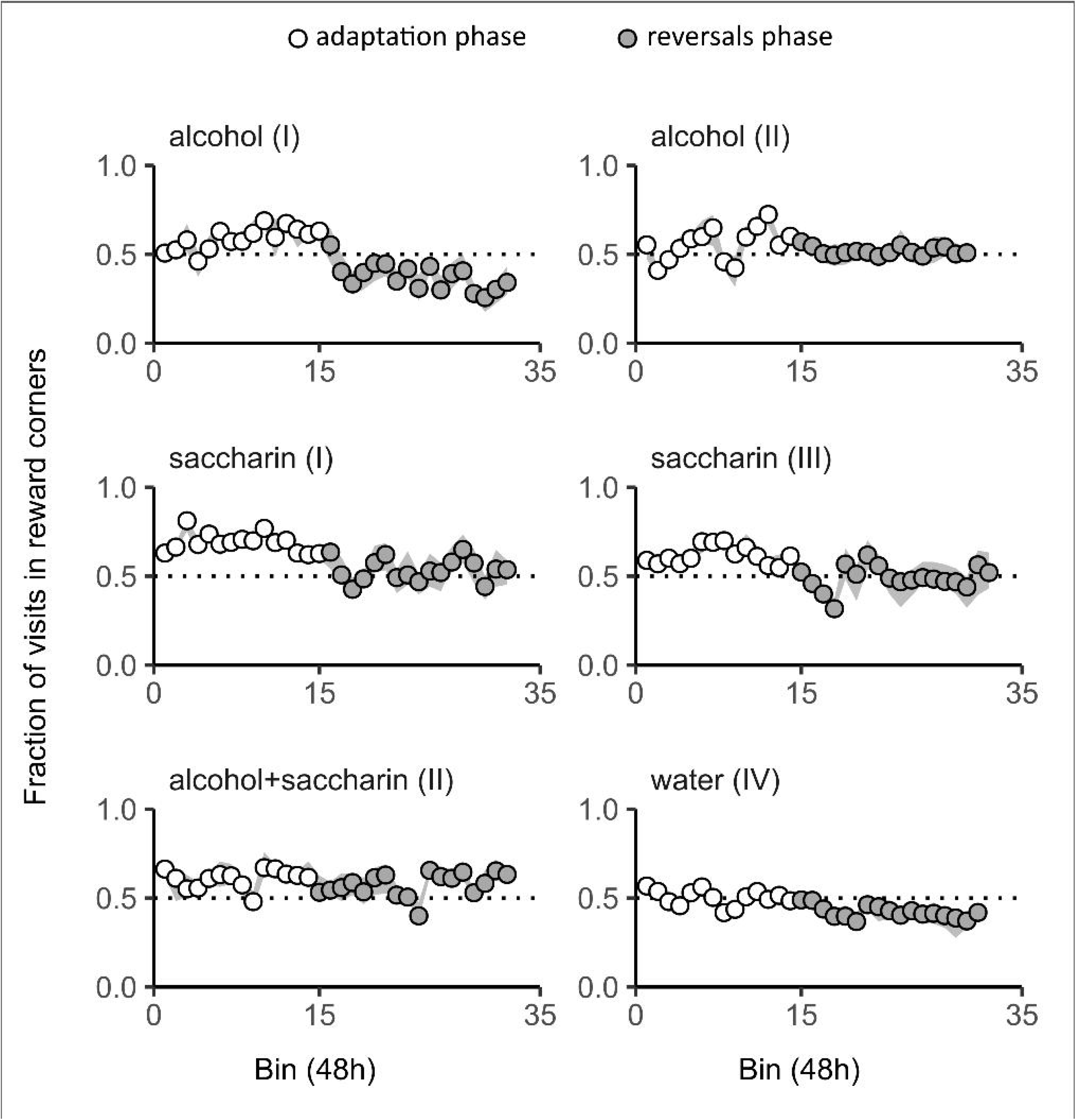
Fraction of the total number of visits corresponding to corners with probabilistic access to the reward per 48h period. White and grey circles correspond to activity during the adaptation and probability reversal stages, respectively. The gray ribbon shows 1st and 3rd quartiles. The type of reward offered is indicated above the graphs. Roman numerals in parentheses indicate the animal cohort.

